# Local adaptation of a fungal pathogen to temperature along a latitudinal gradient

**DOI:** 10.1101/2024.03.04.583296

**Authors:** Quinn N. Fox, Carrie Goodson, Rachel M. Penczykowski

## Abstract

Whether climate warming will increase or decrease prevalence of an infectious disease partly depends on the potential for pathogens to adapt to higher temperatures. This potential can be assessed by investigating regional variation in pathogen thermal performance and testing for local adaptation to current temperature regimes and host populations. We collected seeds of a host plant (*Plantago rugelii*, a perennial herb) and isolated strains of its specialist fungal pathogen (*Golovinomyces sordidus*, a powdery mildew) from five locations along a latitudinal transect from southern Mississippi to northern Wisconsin, USA. In a laboratory experiment, we placed sympatric and allopatric host–pathogen pairings into seven temperature treatments from 7 to 33 °C. We fitted thermal performance curves to pathogen growth data for each strain. Pathogen strains were locally adapted to temperature, with estimated thermal optima ranging from 20.6 °C (southernmost strain) to 16.7 °C (second-northernmost strain) and generally decreasing 0.26 °C for each degree increase in latitude of origin. However, there was no evidence of pathogen local adaptation to sympatric hosts. Given that powdery mildew spores can disperse long distances via wind, our results suggest that northward spread of warm-adapted strains could facilitate pathogen adaptation to warming climates in this and similar systems.

## Introduction

Climate change and infectious disease present simultaneous threats to living organisms worldwide (Chakraborty et al. 2000). Thus, there is an urgent need to understand how host– pathogen systems respond to climate change, including warmer mean temperatures (Harvell et al. 2002, Altizer et al. 2013, Chakraborty 2013, Burdon and Zhan 2020). Climate warming is expected to alter the geographic and seasonal occurrence of many diseases through effects on abundance and physiology of hosts and pathogens (Rohr et al. 2011, Altizer et al. 2013, Claar and Wood 2020, Cohen et al. 2020, Jeger 2022). However, our ability to predict how disease dynamics will change with future warming is limited by our knowledge of the capacity for pathogens to adapt to changing climatic conditions. Moreover, pathogens face selective pressures from hosts as well as from climatic factors. Therefore, it is essential to establish whether pathogen adaptation to climate regimes is robust across standing variation in host resistance.

Within a pathogen species, strains can vary in their thermal tolerances and optima (Kowalski and Bartnik 2010, Voyles et al. 2017). Where temperatures are below optimum for growth of a pathogen on its host, warming should increase disease risk (Cohen et al. 2020). By the same logic, warming should decrease disease risk if temperatures are at or above optimum for pathogen growth on the host (Faticov et al. 2022). Pathogens can evolve to become locally adapted to their abiotic environment if selective pressures act on standing genetic variation to select for individuals most suited to those environmental conditions (Kawecki and Ebert 2004). In some systems, pathogens are adapted to their local temperature regimes (Laine 2008, Stefansson et al. 2013, Korfanty et al. 2023), with thermal optima that correspond to mean temperatures in their local environment (Mariette et al. 2016). In other systems, pathogens have been found not to adapt to increases in temperature (Schampera et al. 2022).

At the same time, many host–pathogen systems exhibit specificity between host and pathogen genotypes. In such systems, strong selection for pathogen infectivity and host resistance causes genetically variable pathogens and their hosts to engage in reciprocal adaptation and counter-adaptation. Given enough time to coevolve, pathogen populations can become locally adapted to their host populations, where infections are more likely in sympatric (local) than allopatric (foreign) pairings (Lively and Jokela 1996, Kawecki and Ebert 2004). Local adaptation is most likely to occur when pathogen populations have high levels of genetic diversity, and thus high potential for novel genotypes that can overcome host resistance (Höckerstedt et al. 2018). High genetic diversity within pathogen populations is typically associated with short generation times, large populations, high mutation rates, and moderate gene flow (Gandon et al. 1996, Höckerstedt et al. 2018). Indeed, many pathogens are found to be more infective on their sympatric than allopatric hosts (Lively and Jokela 1996, Thrall et al. 2002, Kawecki and Ebert 2004, Laine et al. 2011, Koskella 2014). Yet, this is not always the case, even for specialist pathogens that should be under strong selection to infect local hosts (Höckerstedt et al. 2018). Importantly, the performance of pathogen life history traits can depend on three-way interactions among host genotypes, pathogen genotypes, and environmental variables (Wolinska and King 2009, Penczykowski et al. 2016). Therefore, a pathogen that is adapted to local hosts under a given climate regime might not be locally adapted to its hosts under warming.

Here, we tested for local adaptation of a pathogen to thermal regimes and host genotypes along a latitudinal gradient spanning large variation in mean annual temperature. We used an experimentally tractable system comprised of the herbaceous host plant *Plantago rugelii* and strains of its specialist powdery mildew pathogen, *Golovinomyces sordidus,* collected along a latitudinal gradient in North America (Penczykowski and Sieg 2021). We expected that pathogen growth would vary with temperature according to unimodal thermal performance curves, such that a thermal optimum could be estimated for each pathogen strain. We hypothesized that the pathogen would exhibit local adaptation to temperature, where strains from more southern locations would have warmer thermal optima and strains from more northern locations would have cooler thermal optima. Additionally, we hypothesized that pathogen strains would be locally adapted to host populations, with more growth in sympatric than allopatric pairings.

## Materials and Methods

### Study system

Our focal host plant is the rosette-forming herb *Plantago rugelii* Decne. (blackseed plantain). This short-lived perennial grows commonly in pastures and human-disturbed landscapes (e.g., along paths or roads and in mowed, open areas) over much of eastern North America (Penczykowski and Sieg 2021). Our focal pathogen is the powdery mildew fungus *Golovinomyces sordidus* (L. Junell) V.P. Heluta, which only infects plants in the genus *Plantago* (Braun and Cook 2012). Powdery mildews (order Erysiphales) are obligate pathogens that extract nutrients from the epidermal tissue of their hosts. Powdery mildew infections are visually conspicuous due to the chains of asexual spores produced from mycelium on the leaf surface which give infected leaves a characteristic white, dusty appearance. Spores are transmitted passively via wind (Tack and Laine 2014). More than 90% of spores land within two meters of their host plant; however, rare instances of long-distance spore transport allow pathogens to spread over a regional scale (Ovaskainen and Laine 2006, Tack and Laine 2014). Mildews survive winter by producing protective sexual resting structures (chasmothecia) which release ascospores when conditions become favorable in the spring (Tack and Laine 2014).

### Field populations

*Plantago rugelii* seeds and powdery mildew spores used in this study were sampled from field populations spanning a latitudinal gradient of over 1,500 km in the central United States (Fig. 1). We collected *P. rugelii* seeds from healthy-looking plants in each of five populations, listed in order of increasing latitude: (1) McComb, MS, (2) Memphis, TN, (3) St. Louis, MO, (4) Morrison, IL, and (5) Altoona, WI (Fig. 1). Leaves infected with *G. sordidus* were collected from the same populations as the seeds, with the exception of the St. Louis host and pathogen populations, which were sampled 18 km apart. We used climate data from airport weather stations nearest our field populations to characterize average daily temperature from late spring through mid-autumn from 2012-2022 (Fig. 1; data from NOAA and NCEI, 2023).

**Fig. 1.**
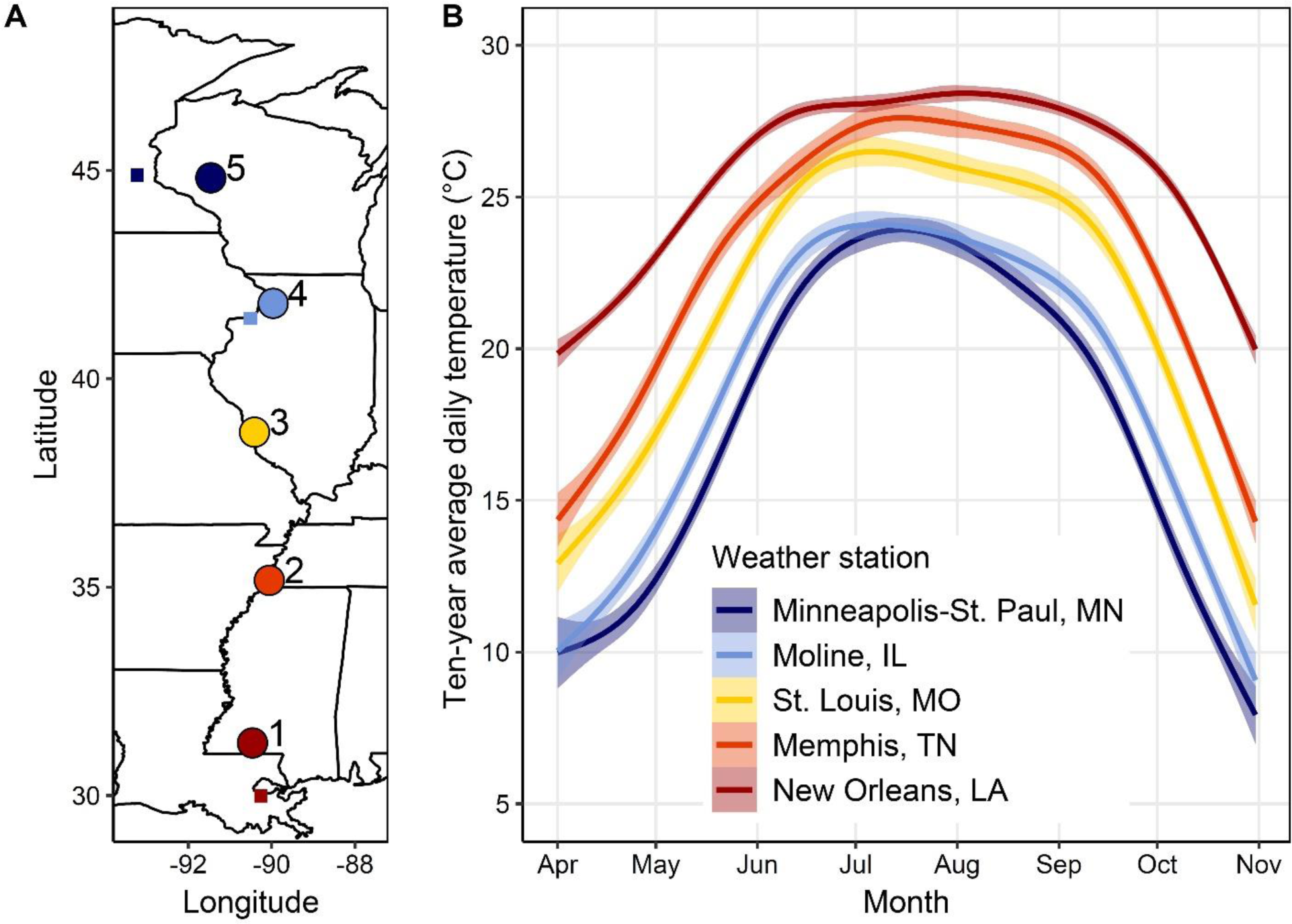
Locations and average temperatures of sampled host and pathogen populations. (A) Populations sampled for *Plantago rugelii* host lines and *Golovinomyces sordidus* pathogen strains used in our experiment, labelled by latitudinal rank. Small squares show locations of airport weather stations, if not located in the same city as the sampled population. (B) Mean (solid line) and 95% confidence interval (ribbon) in average daily temperature between April 1 and October 31 over the 10-year period from 2012-2022 at airport weather stations nearest our sampled populations. Temperature data from NOAA (NOAA and NCEI 2023).

### Plant and mildew preparation

Plants used in this experiment were grown from seeds collected from the five wild *Plantago* populations between 2020-2022. We used a single maternal line per population, where each maternal line originated from a single seed spike from a mother plant. *Plantago rugelii* seeds require cold stratification to increase germination success (C. Goodson, unpublished). Therefore, seeds were placed between layers of wet filter paper in Petri dishes in a 6-7 °C incubator for two weeks in August 2022. Seeds were then moved to a 20 °C incubator for 10 days to allow germination. In a greenhouse, seedlings were initially established in FPX B soil (ProMix), and two weeks later transplanted into individual 4” pots in BM6 All-Purpose soil (Berger). Each pot was covered with an autoclaved, spore-proof pollination bag (PBS International, model 3D.55) to avoid unintentional powdery mildew infection in the greenhouse.

Powdery mildew strains were collected in 2020-2022 by detaching infected leaves from wild host plants using ethanol-sterilized forceps. Infected leaves were placed into Petri dishes containing wet filter paper. Using a sterilized paintbrush, spores from the field-collected leaves were brushed onto healthy leaves detached from greenhouse-grown plants. We isolated pure clonal lines (hereafter, “strains”) by transferring a single chain of asexual spores from an infected to a healthy leaf for three successive generations (∼10 days each; (Nicot et al. 2002). The mildew strains were maintained in growth chambers at 20 °C, 16:8 hours light:dark cycle, and 70% relative humidity (Nicot et al. 2002).

### Experimental setup

Individual leaves were removed from the source plants, cut into 2-cm^2^ fragments and surface-sterilized in 70% ethanol for 20 sec followed by three consecutive 10-sec rinses in autoclaved tap water. The surface-sterilized leaf fragments were placed in Petri dishes, so that each dish contained one leaf piece from each of the five host maternal lines. On November 10th, 2022, each dish was inoculated with a single mildew strain, such that each plate contained one sympatric pairing and four allopatric pairings. To inoculate, we used a single sterilized paintbrush hair to transfer six chains of spores from source mildew lesions onto the center of each leaf piece. The plates of inoculated leaves were placed in growth chambers at each of seven temperature treatments, with three replicate plates of each mildew strain at each temperature (N = 525 leaf pieces). We chose the growth chamber temperatures of 7, 12, 16, 20, 24, 28, and 33 °C to span the typical thermal range for powdery mildew sporulation, where the average thermal minimum (T_min_), thermal maximum (T_max_), and thermal optimum (T_opt_) estimated for several other powdery mildew species are T_min_ = 11.9 °C, T_max_ = 31.2 °C, and T_opt_ = 21.6 °C (Chaloner et al. 2020). Chambers were kept at 16:8 hours light:dark and 60-70% humidity throughout the experiment to promote mildew growth, and plates were spatially randomized within growth chambers every other day.

### Assessment of pathogen performance

Inoculated leaves were examined for progression of asexual growth under a dissecting microscope every other day for 14 days post-inoculation (DPI). Due to a complete lack of mildew growth in the 33 °C treatment, leaves in that treatment were no longer examined after 10 DPI. We quantified mildew growth using a modified five-level categorical index of mildew development (“Bevan score”), where 0 = no growth, 1 = mycelium only, 2 = mycelium and sparse sporulation visible only under a dissecting microscope, 3 = abundant sporulation and lesion size < 0.5 cm^2^, and 4 = abundant sporulation and lesion size > 0.5 cm^2^ (Bevan et al. 1993, Numminen and Laine 2020). The few leaves which had only mycelium (i.e., Bevan score = 1) at 14 DPI were monitored up to 18 DPI for signs of sporulation.

### Statistical analyses

All statistical analyses were performed in R version 4.1.0 (R Core Team 2021). To assess whether the speed of sporulation (i.e., days from inoculation to sporulation) depended on the interaction between host line and pathogen strain, averaged across all temperature treatments, we fitted Cox proportional hazard mixed effects models (function coxme() in ‘coxme’ package; Therneau 2022). We fitted the models to data from each inoculated leaf piece (N = 525), with plate identity included as a random effect because there were five leaf pieces per Petri dish. We compared among a set of models with fixed effects of either host line, pathogen strain, both as main effects, or their interaction. Models were compared using Akaike’s information criterion corrected for small sample sizes (AICc), and the best fitting model was selected using function aictab() from the ‘AICcmodavg’ package (Mazerolle 2020). After fitting these models to data from all temperature treatments combined, we separately fit the models to data from just the coolest three temperature treatments or just the warmest three viable temperature treatments. This was done in order to assess whether the effects of host line, pathogen strain, or their interaction varied qualitatively across the range of temperatures, because models with all seven temperature treatment levels (and their interactions with host line and/or pathogen strain) were over-parameterized and did not converge. We used a similar model comparison approach to evaluate effects of host line, pathogen strain, and their interaction on the final amount of pathogen growth measured as Bevan score at 14 DPI. The Bevan score for each leaf was analyzed as an ordered categorical response variable using cumulative link mixed models (CLMMs) with random effect of plate identity (function clmm() in ‘ordinal’ package, (Christensen 2022)). Separately, we tested for pathogen local (mal-)adaptation to host genotypes as a difference between days to sporulation or final Bevan scores on sympatric vs. allopatric host lines.

For all leaf pieces with some amount of sporulation by 14 DPI (n = 148), we tested for an association between how quickly the pathogen had sporulated (i.e., days from inoculation to sporulation) and the final amount of pathogen growth achieved (i.e., categorical Bevan score at 14 DPI) using a generalized linear model with poisson error distribution. In a mixed effects version of that same model, there was zero variance associated with a random effect of plate.

For each mildew strain, Thermal Performance Curves (TPCs) were fit to the maximum Bevan score (at 14 DPI) among all replicates of all host lines in each temperature treatment. We compared the fits of multiple models commonly used for fitting fungal TPCs (Dumur et al. 1990, Angilletta 2006, Mastrodimos et al. 2019, Omuse et al. 2022). Models were fit using the ‘rTPC’ package pipeline which uses the ‘nls.multstart’ package (Padfield and Matheson 2020, Padfield and O’Sullivan 2021). Models were compared using AIC, and the best fitting model was selected using function aictab() from the ‘AICcmodavg’ package (Mazerolle 2020).

For the best-fitting model, we performed bootstrapping (999 resamplings) to generate 95% confidence intervals for each curve and to estimate average temperature optima (Padfield and O’Sullivan 2021). Then we tested whether thermal optima were significantly negatively related to latitude (i.e., cooler thermal optima for more northern mildew strains), using a bootstrapped linear regression approach (Shocket et al. 2018). We did this by sampling one of the 999 bootstrapped estimates of thermal optima for each of the five mildew strains and fitting a linear regression between those five bootstrapped estimates and their corresponding latitudes of origin. This produced 999 slopes, and we calculated the fraction of those slopes that were less than zero (null hypothesis for this one-sided hypothesis test: > 5% of slopes greater than zero).

## Results

### Time to sporulation

Temperature strongly influenced time to sporulation, with generally faster sporulation at warmer temperatures (up to 24 °C; Fig. 2). Very little sporulation occurred at the lowest (7 °C) or second-highest (28 °C) temperatures, and no sporulation occurred at the highest temperature of 33 °C (Fig. 2). Across all temperature treatments, our model comparisons revealed that host line explained far more variation in time to sporulation than did pathogen strain, and there was no evidence for an interaction between host line and pathogen strain (Table 1). When we fitted the same models to either just the coolest three temperatures (7, 12, and 16 °C; Table S1) or the warmest three viable temperatures (20, 24, and 28 °C; Table S2), the results were consistent, indicating that the lack of interaction between host line and pathogen strain (and lack of pathogen adaptation to local host genotypes) was robust across temperatures. Moreover, there was no significant difference in time to sporulation between sympatric and allopatric host-pathogen combinations (χ^2^ = 0.15, df = 1, p = 0.71). For leaves on which the powdery mildew successfully sporulated, earlier sporulation was significantly associated with greater amounts of growth (i.e., higher Bevan score categories) achieved by 14 DPI (χ^2^ = 59.68, df = 2, p < 0.0001).

**Fig. 2.**
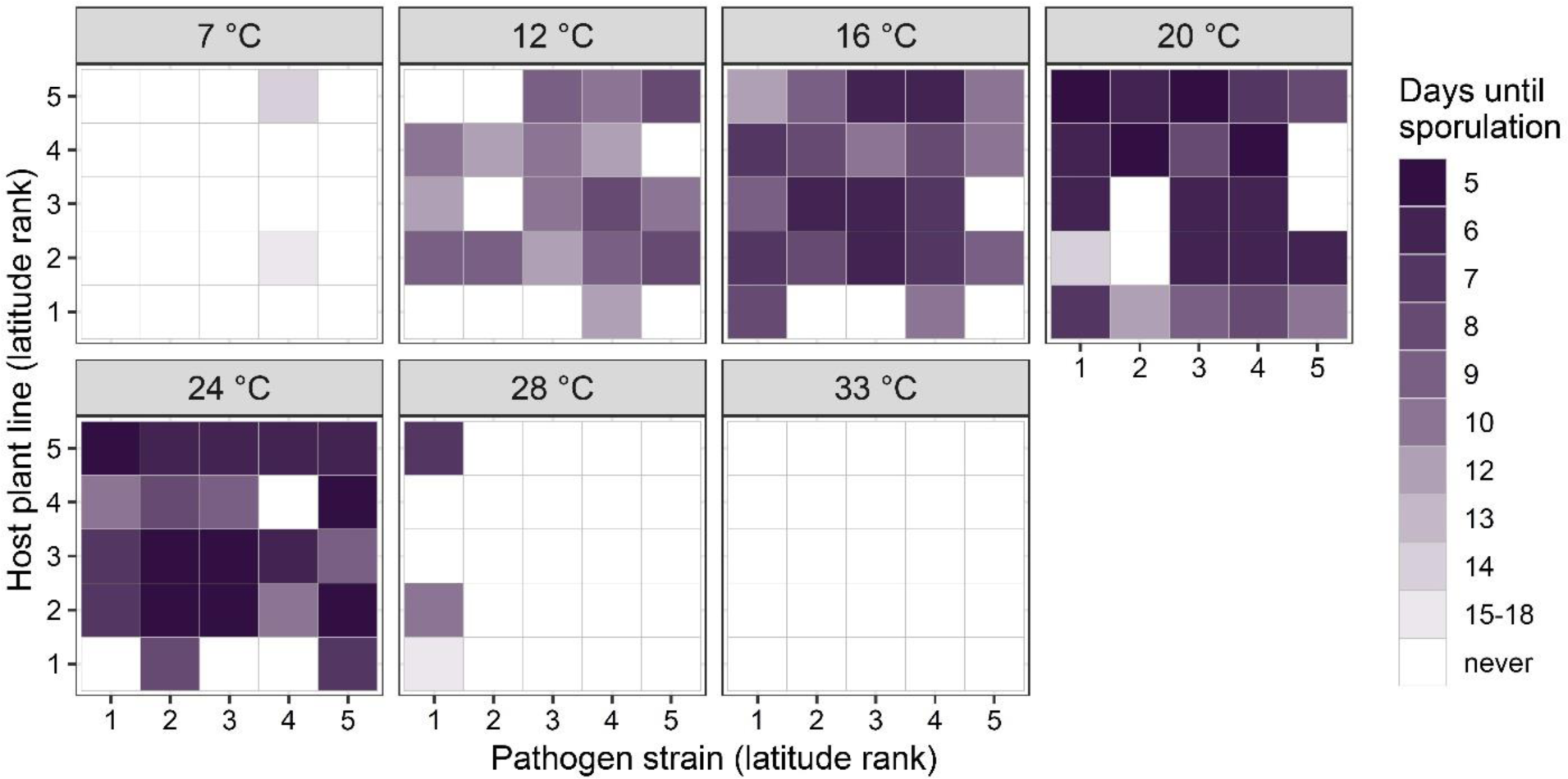
Minimum days until pathogen sporulation for each host–pathogen pairing in each temperature treatment. Sympatric pairs occur along the diagonals of the matrices. The minimum number of days between inoculation and sporulation among three replicate leaves for each host–pathogen–temperature combination is shown here, while each individual leaf is included in the corresponding statistical models (Table 1, Tables S1-S2).

**Table 1.** Comparison of Cox proportional hazards mixed effects models testing effects of host plant line and pathogen strain on days until sporulation, across all temperature treatments. The best performing model is bolded.

| Model terms | k | AICc | ΔAIC | AICc weight | Log-likelihood |
| --- | --- | --- | --- | --- | --- |
| Pathogen strain * Host line | 25 | 1814.15 | 14.38 | 0.00 | -880.77 |
| Pathogen strain + Host line | 9 | 1803.57 | 3.80 | 0.13 | -892.61 |
| Pathogen strain | 5 | 1819.07 | 19.30 | 0.00 | -904.48 |
| **Host line** | **5** | **1799.77** | **0.00** | **0.87** | **-894.83** |

### Final amount of pathogen growth

Temperature strongly influenced the amount of pathogen growth measured as Bevan score. By 14 DPI, all but the southernmost pathogen strain achieved some growth at the lowest temperature (7 °C; Fig. 3). By contrast, only the southernmost strain grew at 28 °C (Fig. 3). No strains grew at 33 °C (Fig. 3). As in our analysis of time to sporulation, across all temperature treatments, the model with only host line performed better than models with pathogen strain or an interaction between host line and pathogen strain (Table 2). The results were consistent when we fitted the models to either the three coolest or three warmest temperature treatments (Tables S3-S4). There was also no significant difference in final pathogen growth between sympatric and allopatric host–pathogen combinations (χ^2^ = 0.86, df = 1, p = 0.35).

**Fig. 3.**
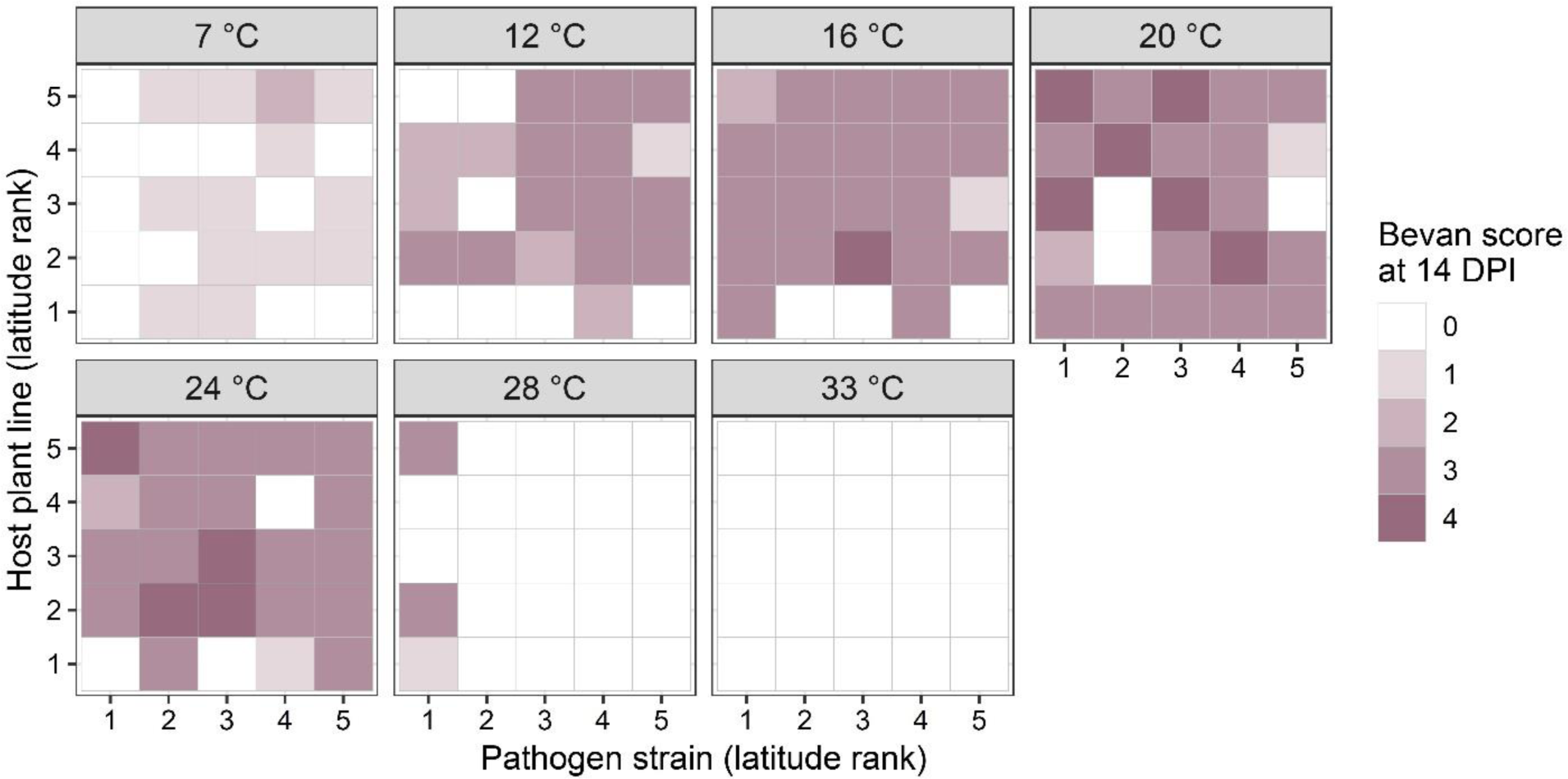
Maximum amount of pathogen growth achieved for each host–pathogen pairing in each temperature treatment. Sympatric pairs occur along the diagonals of the matrices. The maximum Bevan score at 14 DPI among three replicate leaves for each host–pathogen– temperature combination is shown, while each individual leaf is included in the corresponding statistical models (Table 2, Tables S3-S4).

**Table 2.** Comparison of cumulative link mixed models testing effects of host plant line and pathogen strain on the final amount of pathogen growth (Bevan score at 14 DPI), across all temperature treatments. The best performing model is bolded.

| Model terms | k | AICc | ΔAIC | AICc weight | Log-likelihood |
| --- | --- | --- | --- | --- | --- |
| Pathogen strain * Host line | 29 | 1053.50 | 24.76 | 0.00 | -495.99 |
| Pathogen strain + Host line | 13 | 1033.89 | 5.14 | 0.07 | -503.59 |
| Pathogen strain | 9 | 1057.19 | 28.44 | 0.00 | -519.42 |
| **Host line** | **9** | **1028.75** | **0.00** | **0.93** | **-505.20** |

### Thermal optima

The thermal responses of all five pathogen strains were best described by a Gaussian model (Table 5; Fig. 4). The mean bootstrapped estimates of the strains’ thermal optima, listed in order of increasing latitude, were: 20.6 °C (strain 1), 18.4 °C (strain 2), 18.1 °C (strain 3), 16.7 °C (strain 4), and 17.1 °C (strain 5; Fig. 4). These thermal optima were significantly negatively related to latitude (median slope = −0.26; p = 0.001 based on 998/999 bootstrapped linear regressions having negative slopes; Fig. 5).

**Fig. 4.**
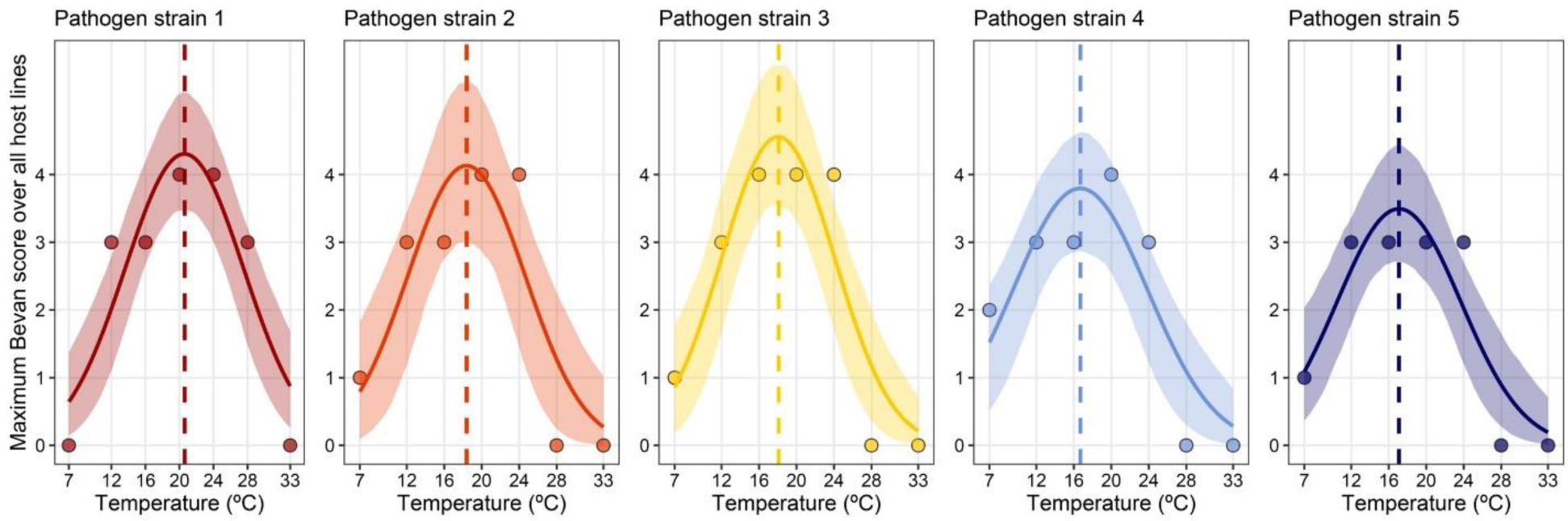
Thermal performance curves for each pathogen strain. Gaussian models (curves) were fit to the maximum values of pathogen growth (points; measured as Bevan score at 14 DPI) obtained across all host plant lines at each of seven temperatures. Pathogen strains are presented in order of latitude rank (1 = southermost, 5 = northernmost). Ribbons give bootstrapped 95% confidence intervals and a dashed vertical line indicates the thermal optimum for each pathogen strain.

**Fig. 5.**
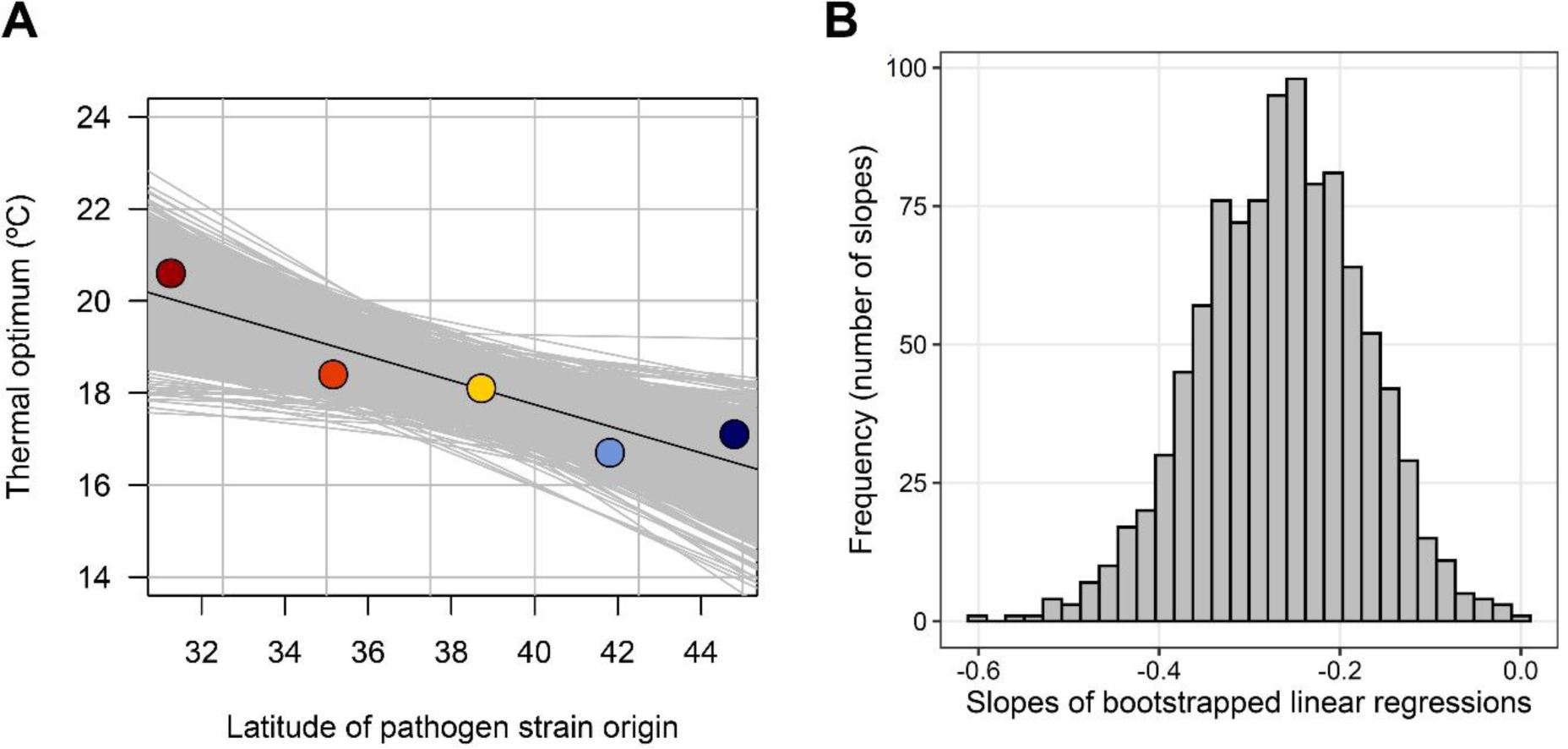
Evidence of pathogen local adaptation to temperature, using latitude as a proxy for local thermal regime. (A) Linear regression (black line) fit to the estimated thermal optimum for each pathogen strain (points color-coded as in Figs. 1 and 4), with 999 bootstrapped linear regressions (gray lines). (B) Histogram of slopes of the 999 bootstrapped linear regressions.

**Table 3.**
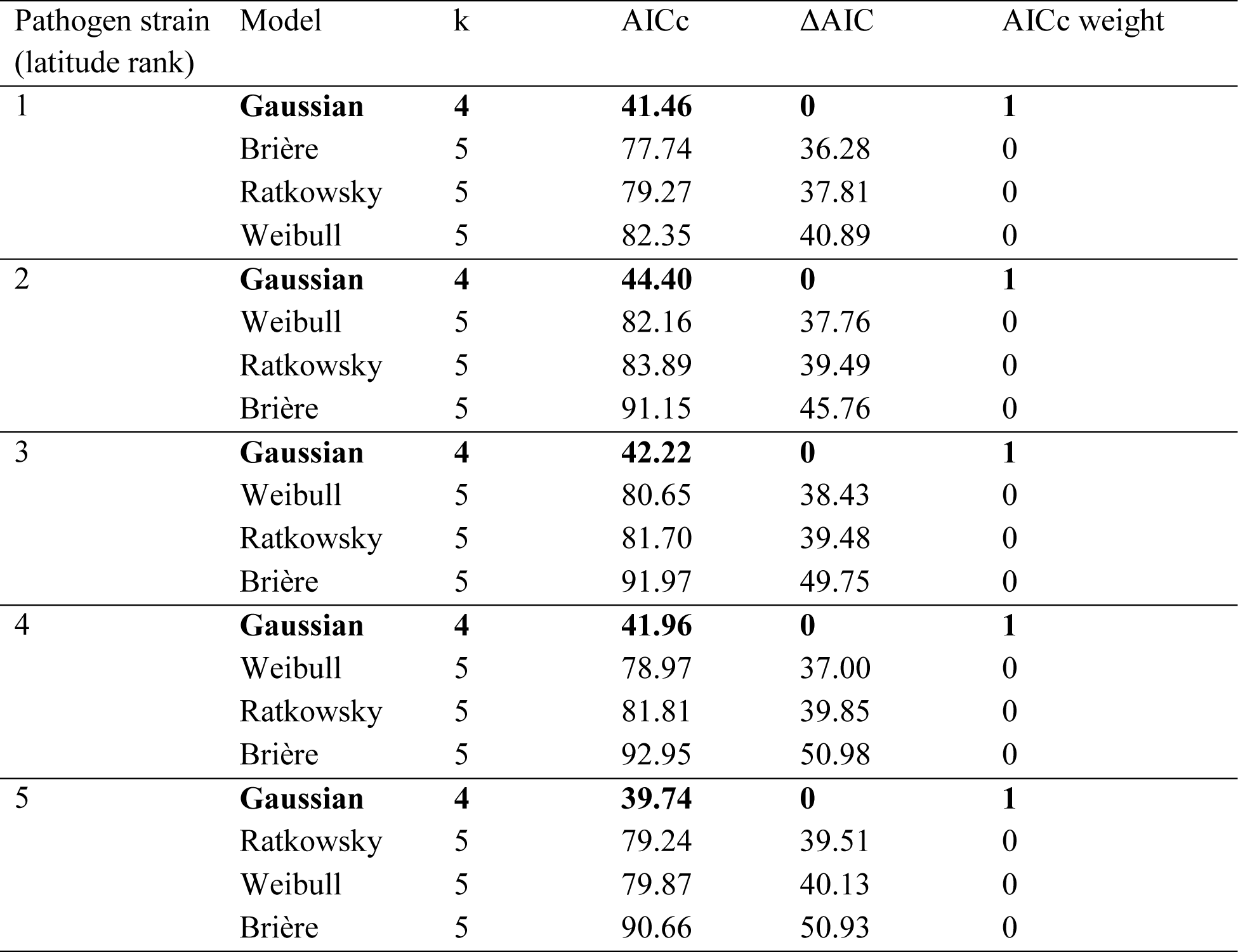
Comparisons of thermal performance curve models fit to maximum growth data from each pathogen strain. The best performing model for each strain is bolded.

## Discussion

Predicting consequences of climate change for infectious diseases remains a formidable challenge in most host–pathogen systems. Even when the many facets of climate change are simplified to focus on warming, there are still complex pathways by which warming can affect hosts, pathogens, and their interaction. Here, we performed a laboratory experiment to test for interactions among host genotypes, pathogen genotypes, and temperatures, using *Plantago* host plants and their powdery mildew pathogen strains collected along a latitudinal gradient spanning wide variation in temperature during the growing season. We applied the temperature treatment to detached leaf pieces immediately after inoculation with the powdery mildew fungus; therefore, our study did not address longer-term temperature effects on growth or susceptibility of the plants themselves. However, we were able to quantify how pathogen strains varied in their thermal performance and to test for local adaptation of pathogen strains to temperature regimes and host genotypes (maternal lines).

As expected, each pathogen strain had a unimodal response to temperature, with growth increasing from cool to intermediate temperatures and sharply declining in the hottest treatments. The temperature range used in our experiment encompassed the upper thermal limits for mildew growth, as no mildew strains grew at 33 °C (and only one grew at 28 °C). Based on thermal constraints documented for other powdery mildew species, we predicted that our lowest temperature treatment of 7 °C would be too low for mildew growth (Chaloner et al. 2020). However, for four of the five mildew strains, we observed some mildew growth at 7 °C, though not necessarily development through to sporulation. Thus, we have less confidence in the ability of our fitted thermal performance curve models to estimate the lower thermal limits for growth of these pathogens. Future experiments with additional colder temperature treatments are needed to assess among-strain variation in lower thermal limits.

In support of our first hypothesis, we found evidence of pathogen local adaptation to regional temperatures. Strains from more southern/warmer populations had higher thermal optima, and strains from more northern/cooler populations had lower thermal optima. There was a 3.9 °C difference between the highest and lowest thermal optima (strains of latitudinal rank #1 and #4, respectively). Notably, the southernmost mildew strain was the only one to successfully grow at the highest survivable temperature (28 °C), and also the only one that failed to establish in the coldest treatment (7 °C). While this pattern might be driven by natural selection for thermal optima closer to average temperatures, it is also possible that the observed differences in thermal performance are the legacy of other neutral or adaptive processes. Future analysis of population genetic structure and genotype-environment associations will be helpful for resolving the causes of the observed pattern of local adaptation.

Contrary to our second hypothesis, there was no evidence of local adaptation of pathogen strains to their sympatric host lines. The best models for both time to sporulation and final pathogen growth did not include interactions between host line and pathogen strain, whether these models were fit to the coolest three temperatures, warmest three viable temperatures, or data from all temperature treatments combined. The lack of a pattern of local adaptation is not entirely surprising, given that host–pathogen coevolution is a dynamic process, and a given pathogen population could be locally adapted to their hosts in some years but not others (e.g., due to parasite-mediated selection for host resistance or stochastic overwintering or dispersal events). Moreover, due to constraints on mildew and *Plantago* seed availability, we used host lines and pathogen strains that had been collected from field populations over the span of a few years (i.e., our sympatric host and mildew pairings were not always exactly contemporary).

Additionally, in a review of studies examining parasite local adaptation, the authors found that over half of the case studies did not demonstrate local (mal)adaptation (Greischar and Koskella 2007). However, the lack of overall interactions between host lines and pathogen strains (regardless of sympatric/allopatric status) is surprising, given that powdery mildews tend to show genotypic specificity in their interaction with a congeneric host, *Plantago lanceolata*, in its native range of Europe (Laine 2004). Further research on genotypic and phenotypic variation of *Plantago rugelii* in eastern North America (where it is endemic) will be necessary for explaining the apparent lack of specificity observed here.

Plant disease epidemics have the potential to cause significant ecological, agricultural, and economic damage, and are considered one of the foremost challenges to achieving global food security (Velásquez et al. 2018). The environmental conditions in which a host and its pathogens interact has a significant impact on the outcome of the relationship. Understanding and predicting impacts of climate change on plant–pathogen interactions is therefore a huge societal challenge. Here, we documented among-strain variability in thermal performance of a wild plant pathogen over a regional scale and a pattern of pathogen local adaptation to temperature. For pathogens with high potential for dispersal––including wind-dispersed fungi such as powdery mildews––this existing regional variation in thermal performance could facilitate adaptation to future warming.

## Acknowledgements

We thank Cheyenne Morris, Lily Goldberg, and Allison Rea for assistance with lab and greenhouse work. Mike Dyer provided greenhouse support. We thank Marta Shocket for advice on statistical analysis of bootstrapped parameter estimates. This material is based upon work supported by the National Science Foundation under Grant No. DEB-2304479 awarded to RMP.

## Author Contributions

RMP and QNF conceived of study design. CG, RMP, and QNF collected plant and pathogen materials from field populations. CG prepared host and pathogen materials for the experiment. QNF performed experiment. QNF and RMP performed statistical analyses. QNF and RMP wrote the first draft, and all authors contributed to subsequent drafts of the manuscript.

---

**Table S1.** Comparison of Cox proportional hazards mixed effects models testing effects of host plant line and pathogen strain on days until sporulation, across the coolest three temperature treatments (7, 12, and 16 °C). The best performing model is bolded.

| Model terms | k | AICc | ΔAIC | AICc weight | Log-likelihood |
| --- | --- | --- | --- | --- | --- |
| Pathogen strain * Host line | 25 | 741.99 | 14.77 | 0.00 | -342.73 |
| Pathogen strain + Host line | 9 | 730.96 | 3.74 | 0.13 | -356.06 |
| Pathogen strain | 5 | 755.83 | 28.60 | 0.00 | -372.78 |
| **Host line** | **5** | **727.22** | **0.00** | **0.87** | **-358.48** |

**Table S2.** Comparison of Cox proportional hazards mixed effects models testing effects of host plant line and pathogen strain on days until sporulation, across the warmest three temperature treatments at which powdery mildew grew (20, 24, and 28 °C). The best performing model is bolded.

| Model terms | k | AICc | ΔAIC | AICc weight | Log-likelihood |
| --- | --- | --- | --- | --- | --- |
| Pathogen strain * Host line | 25 | 834.99 | 28.38 | 0.00 | -389.23 |
| Pathogen strain + Host line | 9 | 813.22 | 6.61 | 0.04 | -397.19 |
| Pathogen strain | 5 | 820.12 | 13.51 | 0.00 | -404.92 |
| **Host line** | **5** | **806.61** | **0.00** | **0.96** | **-398.17** |

**Table S3.** Comparison of cumulative link mixed models testing effects of host plant line and pathogen strain on the final amount of pathogen growth (Bevan score at 14 DPI), across the coolest three temperature treatments (7, 12, and 16 °C). The best performing model is bolded.

| Model terms | k | AICc | ΔAIC | AICc weight | Log-likelihood |
| --- | --- | --- | --- | --- | --- |
| Pathogen strain * Host line | 29 | 516.61 | 22.20 | 0.00 | -224.84 |
| Pathogen strain + Host line | 13 | 498.17 | 3.76 | 0.13 | -235.22 |
| Pathogen strain | 9 | 523.54 | 29.12 | 0.00 | -252.35 |
| **Host line** | **9** | **494.41** | **0.00** | **0.87** | **-237.79** |

**Table S4.** Comparison of cumulative link mixed models testing effects of host plant line and pathogen strain on the final amount of pathogen growth (Bevan score at 14 DPI), across the warmest three temperature treatments at which powdery mildew grew (20, 24, and 28 °C). The best performing model is bolded.

| Model terms | k | AICc | ΔAIC | AICc weight | Log-likelihood |
| --- | --- | --- | --- | --- | --- |
| Pathogen strain * Host line | 29 | 498.36 | 32.91 | 0.00 | -215.72 |
| Pathogen strain + Host line | 13 | 471.70 | 6.26 | 0.04 | -221.99 |
| Pathogen strain | 9 | 480.51 | 15.07 | 0.00 | -230.84 |
| **Host line** | **9** | **465.44** | **0.00** | **0.96** | **-223.30** |

